## Supplementary figures and images for "Corticospinal excitability remains unchanged in the presence of residual force enhancement and does not contribute to increased torque production"

### Supplemental Figure 1

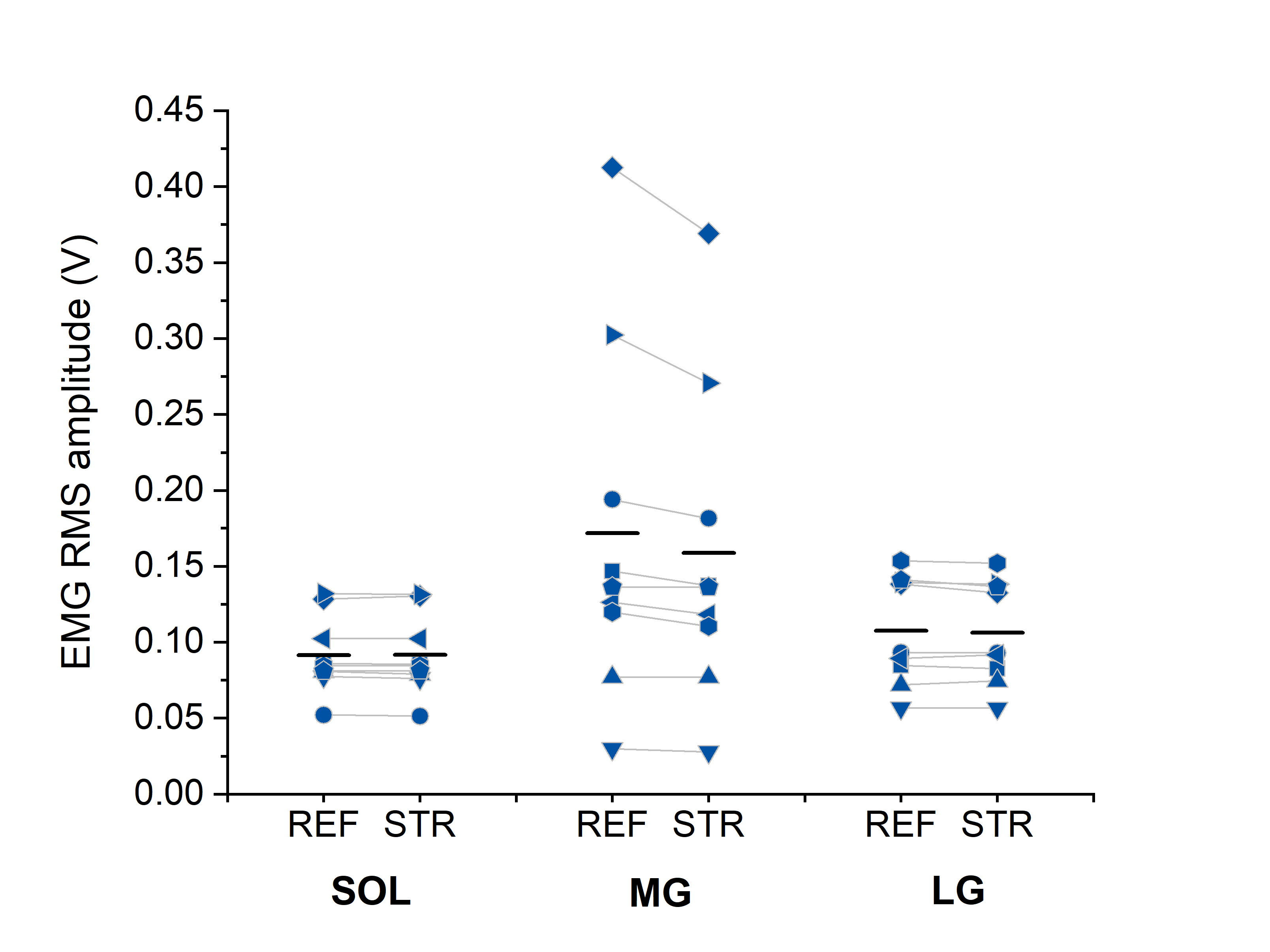
